# Pan-cortical area sensorimotor network coordination during motor learning of forelimb-reaching task in the marmoset

**DOI:** 10.64898/2026.02.23.706473

**Authors:** Yukako Yamane, Teppei Ebina, Masanori Matsuzaki, Kenji Doya

## Abstract

Motor learning alters activities in multiple brain areas. While learning-induced activity change in these areas has been investigated, how the information flow changes in the network across those areas remains to be thoroughly examined. We analysed wide-field calcium imaging data spanning from the premotor cortex to the parietal cortex of marmosets while they learned a two-target forelimb-reaching task. We applied non-negative matrix factorization (NMF) to the activity data and extracted about 30 localized activity components. Encoding model analysis indicated that learning was associated with a decrease in activity components related to hand movements, and an increase in those related to external and reward signals. Causality analysis by embedding entropy (EE) revealed increases in causal links across activity components in different areas and stabilization of the network structure with behavioural improvements. These results indicate that motor learning entails both a redistribution of task-related activity and a reorganization of large-scale cortical network interactions.

## INTRODUCTION

The regions involved in motor learning include multiple cortical and subcortical areas. In the primary motor area, synaptic plasticity ^46,47^ and reduced activity variability^35^ during learning are reported. The dorsorostral and dorsocaudal premotor areas (PMdr, PMdc) show changes of selectivity across learning^11^. As motor skill acquisition involves changes in perceptual processing, the somatosensory area is also involved in motor learning^33^. Disruption of somatosensory sensation^7^ or activity^28^ prevents learning. The connection between the somatosensory and motor areas changes with learning^19^. In addition, the parieto-frontal areas have been posited as crucial for sensorimotor transformation^4^. The parietal area provides strong inputs to the motor areas^2, 25^, and the rostral posterior parietal cortex has maps with similar organization to the movement domain maps of M1 and the premotor areas^21^, which indicates a functional relationship.

Multiple brain areas involved in motor learning should change their causal relationships across learning. Previous studies demonstrated network changes in rodents^26,15^. Even though subdivisions of the primate frontal and posterior parietal cortex are involved in higher-order cognition and sensorimotor integration, they do not exist as separate areas in rodents^20,25^. In order to understand motor learning in primate including humans, investigation of changes in the causal relationships between multiple areas including higher order sensorimotor areas is essential.

The marmoset has a brain architecture similar to that of macaques^34,2^, whereas nearly non-existent cortical sulci and flat structure is well-suited for imaging studies. Successful imaging recordings have been obtained from awake marmosets^22,11,12^. Findings from marmosets can be translated more easily to macaques and human brain functions compared to findings from rodents.

We analysed wide-field calcium imaging data of marmosets previously reported ^11^, spanning the premotor to parietal cortices during learning of a two-target forelimb-reaching task. We extracted several tens of local activity components by non-negative matrix factorization^24^ (NMF) and examined their dependence on behavioral parameters, as well as causal relationships using embedding entropy^42^ (EE).

## RESULTS

### Animal behaviour during the learning of a two-target-reaching task

Three marmosets (mo1 (female), mo2 (male) and mo3 (male)) were trained for 4–12 weeks to perform a target-reaching task that involved pulling the 2-dimensional manipulandum to match the visual cursor on the monitor to the target^11^. Thereafter, they began to learn the two-target-reaching task (Figure 1A) that involved pulling (down) or pushing (up) their manipulandum, and we focused on the learning process (3-6 weeks). Figure 1B shows the cursor trajectory during the reaching time in the early and late sessions. Since the animal had already learned the pulling (down) trials, error trials mostly extended downwards in the early session whereas, in the later sessions, the trajectories showed relatively less variability and were straighter. Figure 1C shows the mean and standard deviation of vertical trajectories triggered by motion onset. The standard deviation decreases across session in up trials. To evaluate the decrease of the variability of cursor trajectories, we calculated intrasession deviations from the mean trajectory and compared them across sessions (Supplementary Figure S1A). In pushing trials, the trajectory deviations decreased (compared to bluish trials, reddish traces show lower values). However, in pulling trials, the trajectory deviations did not decrease across sessions, except for mo3, and was around 0.3 s after motion onset which, in most trials, was the reward-delivery timepoint for mo3. Regardless of the target position, mo3 tended to move the manipulandum largely and sometimes continued this even after reward delivery and showed a different strategy for learning from that of the other two monkeys. Furthermore, there were some interindividual differences for behaviour in reaction time that, in successful trials, was longer in earlier sessions in up-target trials (Figure 1D, upper panel) than in down-target trials (lower panel) in mo1. However, in both the up and down trials, mo2 showed a decrease in reaction time that indicated confusion about behaviour in both directions in the early session. In the up trials, mo3 showed a mild improvement in reaction time across sessions. Despite interindividual variability in learning strategies, the trajectory variability and reaction time decreased in the up trials, indicating increased efficiency through behavioural changes across sessions. The success rates were relatively high across sessions and the details of the reason for that is described in Supplementary information figure S1.

**Figure 1.**
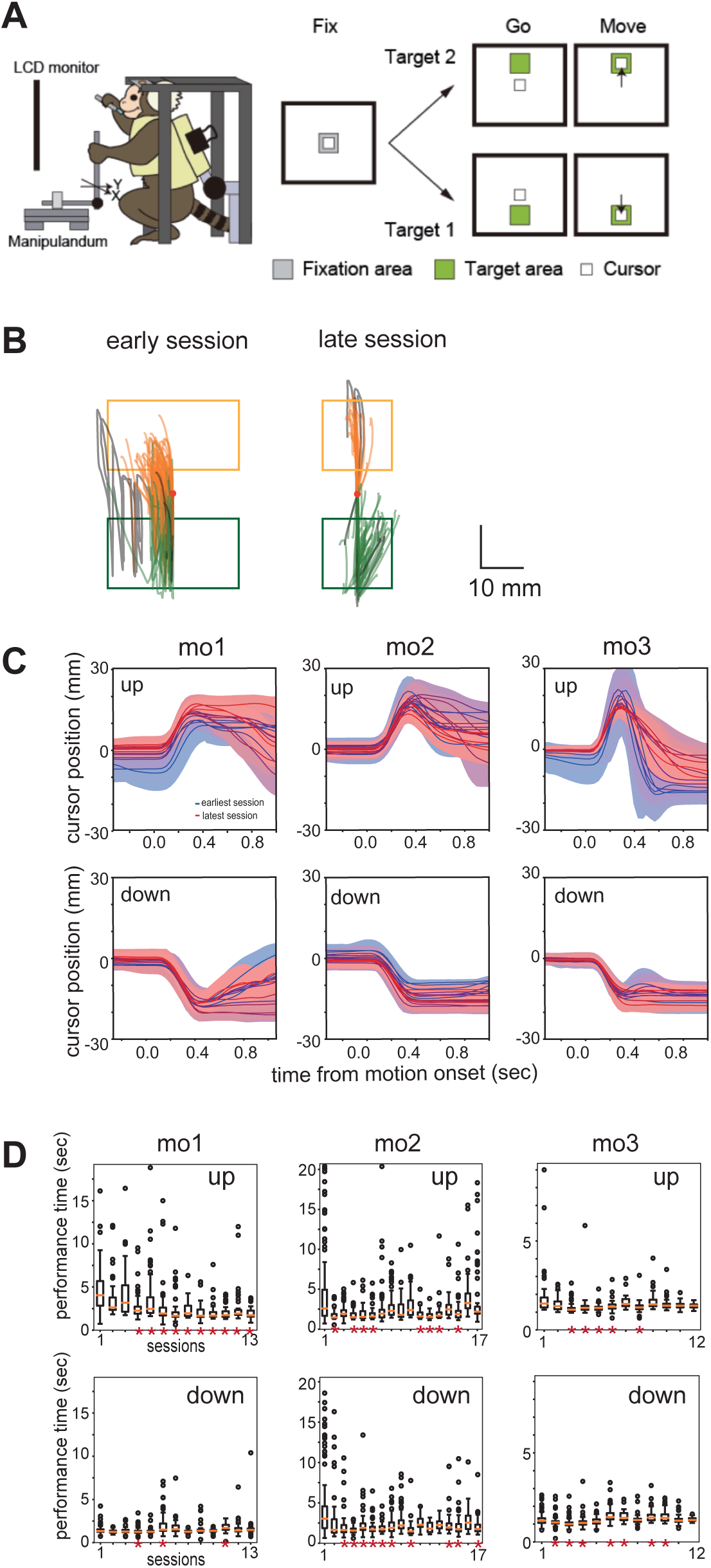
Task description and performance across learning sessions A: Schematics of the two-target reaching task. B: Trajectory of the cursor in earlier and later sessions for monkeys during the reaching time (108 and 111 trials for early and late sessions, respectively). The colour corresponds to the target position (orange: up success, green: down success, and grey: failure trials). The red dots correspond to the centre of the monitor screen. The rectangles represent the accepted areas inside the target. Note that the rectangular size was larger in the early sessions to make the task easy and motivate the animal to continue learning. C: Mean vertical trajectory triggered by motion onset in each session. Colours correspond to sessions. The coloured areas correspond to the standard deviation. D: Reaction times across sessions in the up (upper panels) and down trials (lower panels). The horizontal orange lines are the medians, the upper and lower borders of the boxes are the 25th–75th percentiles, and the upper and lower whiskers represent the maximum and minimum values of the non-outliers, respectively. Dots are outliers. Red asterisks on the x-axis indicate a significant difference between the first session and the other sessions (dscf-test, p<0.05).

### Non-negative matrix factorization for dimension reduction of imaging data

During a two-target reaching task, wide-field calcium imaging was performed with GCaMP6s at 30Hz frame rate. The imaging field-of-view (12 to 14 mm length) included parts of the premotor, motor, somatosensory, and parietal areas (Figure 2A). We abbreviate premotor, motor, somatosensory, and parietal areas as F, M, S, and P. To extract similar activity pattern and reduce the dimensionality of the data, we performed non-negative matrix factorization^24^ on the Δ F/F movies. We concatenated the data of all sessions after correcting the shifts in the field of view (See METHODS) and extracted common components across sessions (approximately 30,000 pixels common across all sessions; Figure 2B). We tested NMF with different numbers of components and set the number of components to be 50 (METHODS and Supplementary information S2-1A). NMF of the concatenated data yielded 50 footprints (*W*) and 50 corresponding time-courses (*H*). After excluding footprints that were strongly correlated with the blood-vessel patterns (Supplementary information S2-1B), 34, 33, and 29 activity components remained for mo1, 2, and 3, respectively. Majorities of the footprints were spatially localized within one of four areas (Supplementary information S2-2). Figure 2C shows the footprints by contour lines (50% of footprint peak) and Figure 2D shows some examples of their time courses. The footprints of activity component were distributed throughout the ROI with only small overlaps of up to 2 to 3 components (Supplementary information S2-3).

**Figure 2.**
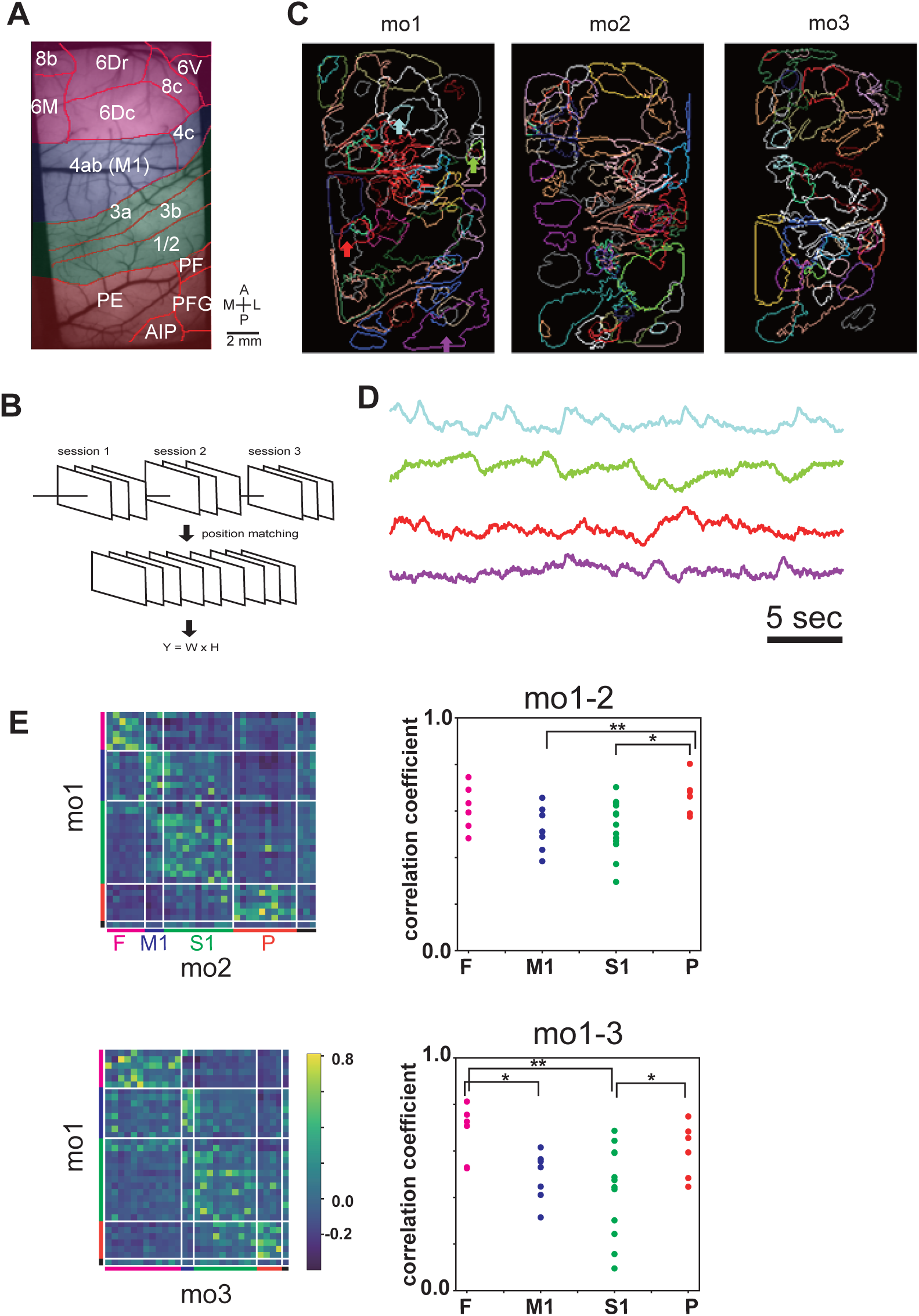
Dimension reduction by NMF A: Field of view of mo1.The red lines represent the putative boundary lines of the cortical areas determined using ICMS. Transparent colors indicate the difference of 4 areas: F(magenta), M1(blue), S1(green), and P(red). B: Schematics of the data preparation for NMF. C: Footprints of all extracted components. The colours correspond to the individual components, and their contours represent 50% of the peak weights of the individual components. The arrows indicate the position of the example components in D. D: Example time course of components of mo1. E: The similarity of footprint patterns between mo1 and mo2, or mo1 and mo3, evaluated as correlation coefficients of pixel values. All pairs of correlation coefficients (Left) are ordered by the categorized four areas (F, M1, S1, and P) and an uncategorized area (black). The correlation coefficients of component combinations showing the best match were plotted (Right). *p<0.05, **p<0.01, Mann–Whitney U test; n=6,8,13 and 6 for F, M1, S1 and P, respectively.

### Similarity of the component-footprint pattern among the subjects

To examine the consistency of activity-component footprints among the subjects, we calculated Pearson’s correlation coefficients between the footprints of one monkey with another for all component pairs (Figure 2E left). The best-matched activity components were predominantly found in the same cortical areas (Figure 2E right). The correlation of footprints across animals within motor and somatosensory areas were lower than those within premotor and parietal areas (Figure 2E right). The higher inter-subject variability in motor and somatosensory areas is possibly attributable to variations in detailed body movements or individual differences of detailed cortical maps.

### The time course of activity components

Figure 3 illustrates how each activity component is modulated around cursor movements using event-triggered averaging. Even among activity components with highly similar footprints, their temporal profiles in triggered averaging were not necessarily alike. For example, activity component F1 showed a high footprint correlation across monkeys (r = 0.75 and 0.81 for mo1-mo2 and mo1-mo3; Fig. 3A), but their temporal profiles of event-triggered average did not closely resemble each other. For example, in mo1F1 and mo2F1, activity was modulated in opposite directions for up and down movement but this contrast was not prominent in mo3F1 (Fig3B).

**Figure 3.**
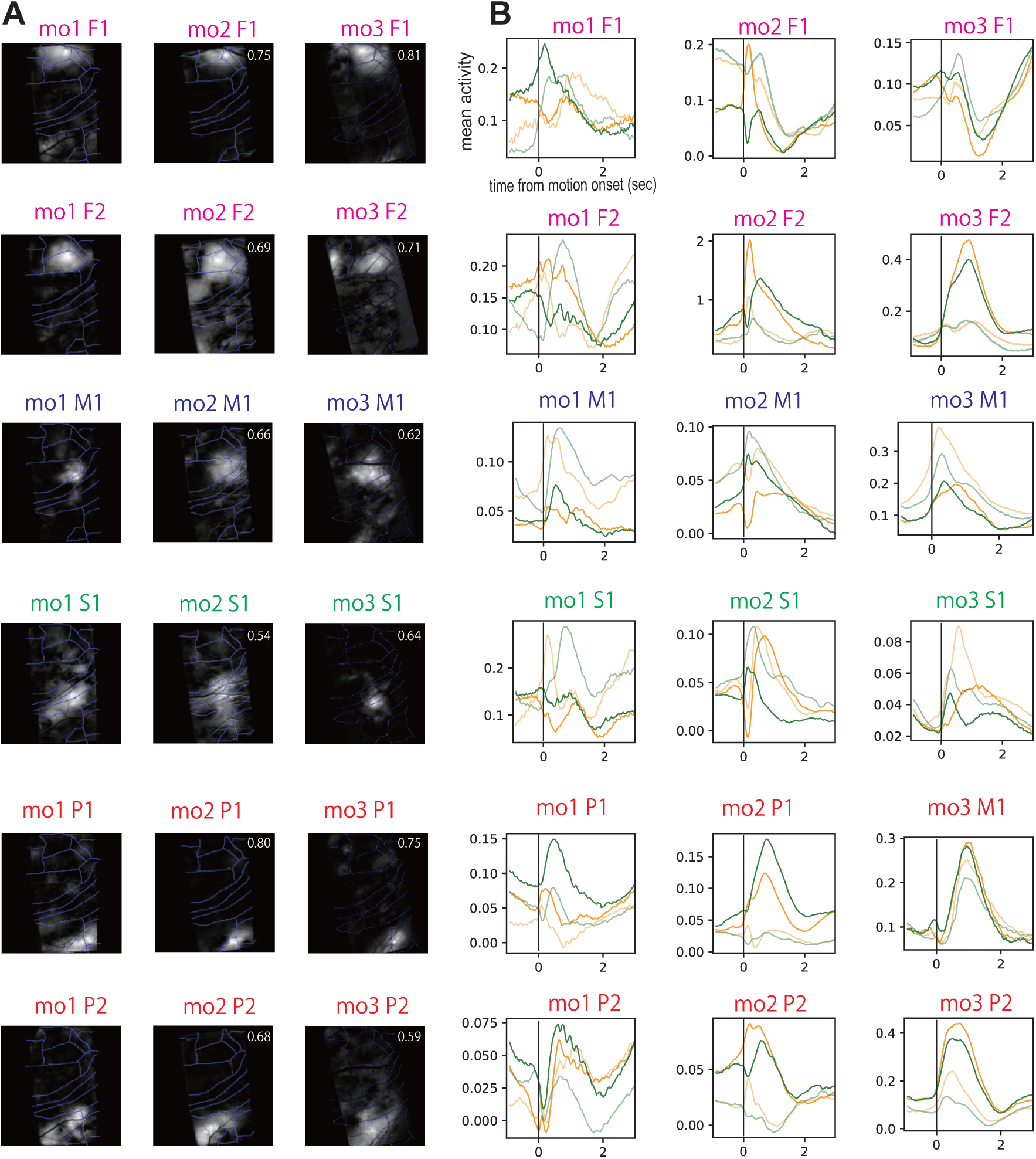
Comparison of activity component footprints and time courses. A: Example footprints of activity components that showed high correlation across animals. White numbers indicate the correlation coefficients between the footprints of mo1 and mo2 or mo3. B: Cursor-movement-triggered averages for up (orange) and down (green) trials. Translucent lines represent averages from early sessions, and opaque lines represent averages from later sessions.

Triggered averaging revealed diverse patterns of learning-related changes, including decreased amplitudes for both up and down movements (e.g., mo1M1), increased amplitudes (e.g., mo3F2), and changes in waveform shape (e.g., mo1F2).

### Linear encoding model of activity components

To investigate the relationship between behaviour and neural activity component time-courses, we created linear encoding models for each activity component in individual sessions. The independent variables were: trial start timing, target (up/down), vertical cursor position, cursor speed, error of the cursor from the target, rewarded timing, unrewarded timing, and mouth-opening angle. As neural activity may precede or follow changes in behavioural parameters, we included 21 Gaussian kernels with lags for each independent variable, which yielded 189 parameters (nine behavioural parameters and 21 lags) for each model. For fitting, we used LASSO to keep the number of effective parameters low (details of the model and their selection in the METHODS). Figure 4A-C shows two representative set of models of activity components of mo1 that showed relatively high scores (R^2^: variance explained) (Supplementary information S4-1A) with positive lags, which indicates that cortical activity precedes mouth movement or cursor movement (Figure 4A, B). Figure 4C shows the actual and predicted time-courses of the model for one session, indicating that the model accurately predicts the time course. The mean R^2^ score of all models across activity components and sessions were 0.38, 0.35, and 0.39 for mo1, 2, and 3, respectively (Supplementary information S4-1B). Thus, the included parameters alone cannot fully explain the cortical activity during task performance.

**Figure 4.**
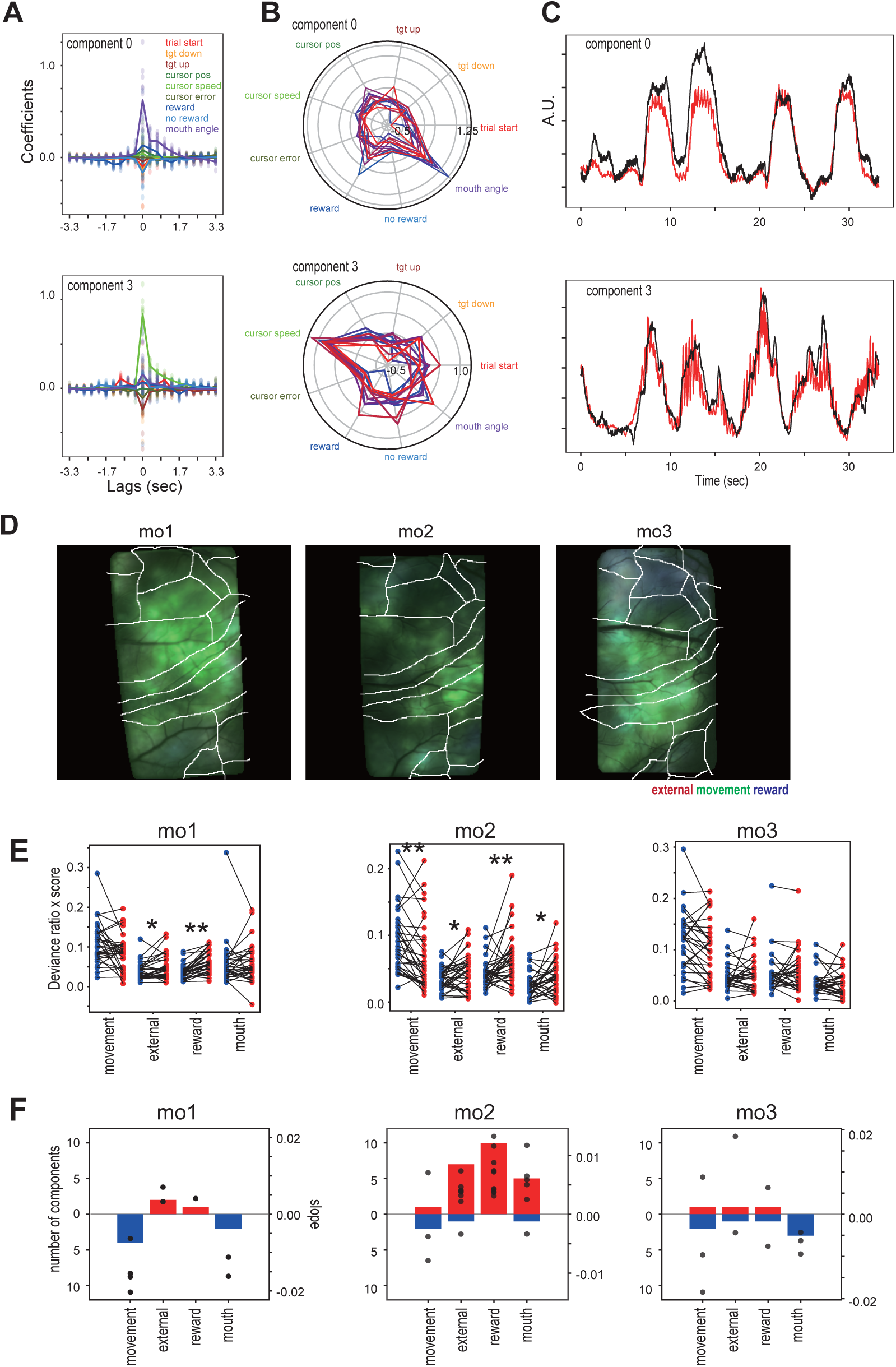
A: The upper and lower panel shows coefficients (β) of the model across lags of different example activity components. Colour indicates the type of exogenous nine variables. Each point corresponds to the coefficient of the model of a session. The lines are mean across sessions. B: The summed coefficients across lags are shown for each corresponding exogenous variables. Colour corresponds to sessions. C: Predicted (red) and actual (black) time courses of the example activity components. D: Deviance ratio for each parameter category projected onto the footprint. Three different categories (external, movement, and reward) of deviance ratio are assigned to red, green, and blue colour, respectively. E: The comparison of mean sDR across early sessions (blue dots) and across later sessions (red dots) for individual activity components. F: The right axis and black dots indicate the slope of the sDR. The left axis and coloured bars indicate the number of models with significant slopes. Only the activity component with a significant slope is shown (p<0.05, Wald’s test).

The R^2^ scores were consistent across sessions, especially for the high-scoring activity components (Supplementary information S4-1C, horizontal striped patterns). Regression coefficients, beta were also consistent across sessions. Supplementary information S4-1D shows Spearman’s correlation coefficients of the beta-coefficients of consecutive sessions. Relatively high consistency across consecutive sessions indicates similarity of model between sessions.

Next, we checked the peak lag time (largest beta-coefficient) for each behavioral variable (Supplementary information S4-2). The peak lag time for self-movement (cursor position, cursor speed, mouth angle) showed strong peak around 0 lag or earlier (neural activity is earlier than the behaviours). The peak lag time for the external parameters were later. These observations are consistent with the general understanding of early motor activity before movement and response latency for sensory response.

To visualize the difference in effective parameters over cortical areas, we calculated the deviance ratio, which is the reduction of the reduced model in the log-likelihood from the full model, scaled by the difference between the log-likelihood of the full and null models. The full model included all five behavioural parameters. The four reduced models did not include external parameters (trial start timing and target up and down timing), movement-related parameters (cursor position, cursor speed, and error of the cursor from the target), reward-related parameters (rewarded timing and unrewarded timing), or mouth-movement-related parameter (mouth angle). Thus, the deviance ratio evaluates the importance of behavioural parameters for explaining the activity component. Figure 4D shows the mean deviance ratios across sessions projected onto the cortical footprints for the three parameter categories (external, movement and reward related parameters). All animal showed a relatively high deviance ratio for the movement related parameters (indicated by green colour) and premotor and parietal areas showed relatively high deviance ratio for reward related parameters (indicated by blue colour). On the other hand, deviance ratio for the external parameters were not high (absence of red colour).

### Change of the model property across sessions

We then inspected the change of model property across sessions by evaluating the score-weighted deviance ratio (sDR) across sessions. Figure 4E shows the mean sDR values across early (blue dots) and later (red dots) sessions for individual activity components. For movement-related components, sDR tended to decrease in later sessions (mo1: early = 0.11, late = 0.087, p = 0.19; mo2: early = 0.092, late = 0.069, p = 0.001; mo3: early = 0.12, late = 0.10, p = 0.056). In contrast, both external- and reward-related components showed increased sDR in later sessions. For external-related components: mo1: early = 0.040, late = 0.049, p = 0.020; mo2: early = 0.035, late = 0.045, p = 0.044; mo3: early = 0.048, late = 0.052, p = 0.97. For reward-related components: mo1: early = 0.037, late = 0.063, p = 2.7 × 10⁻⁶; mo2: early = 0.043, late = 0.065, p = 0.0023; mo3: early = 0.056, late = 0.058, p = 0.37. We linearly fit the change of this value for four different behavioural categories across sessions and found a tendency of decreasing in movement-related values and increasing of external and reward related values (Figures 4F). The result indicates a change in the component activity related to one’s own movement to that related to learned behaviour (external or reward).

To summarize the results of linear modelling, some activity as had high regression scores, the score values and beta-coefficients were relatively stable across sessions, indicating relatively constant results. Areas that showed high dependency on movement-related parameters were widespread across many areas in the ROI. Despite the limited explained variance, the model showed that the component activity changed from that related to one’s own behaviour to that related to learned behaviour across sessions.

### Relationship between activity components evaluated by embedding entropy analysis

We examined the interrelationship among the activity components. Embedding entropy^42^ (EE) is used to examine causal relationships among time-series, such as Granger causality; however, it applies to nonlinear dynamics. EE evaluates the relationship by predicting a time course based on another time course projected onto the delay-embedding space (See METHODS). We applied the time-series of each activity component pair in each session. Figure 5A shows the mean EE values across sessions for each monkey. A high value indicated strong predictability (causal relationships). Matrix is generally symmetric, indicating that the predictions in both directions are relatively similar. This observation is reasonable, considering the high rate of reciprocal connections in the brain. Areas with high EE values were distributed, and we did not find a particular area in which all activity components had high or low EE values; however, there was some variation within individual areas. We also examined whether there is interaction between EE values and the dominant explaining variable category identified in linear regression. We did not find any strong relationships consistent across monkeys (Figure 5A vertical bars).

**Figure 5.**
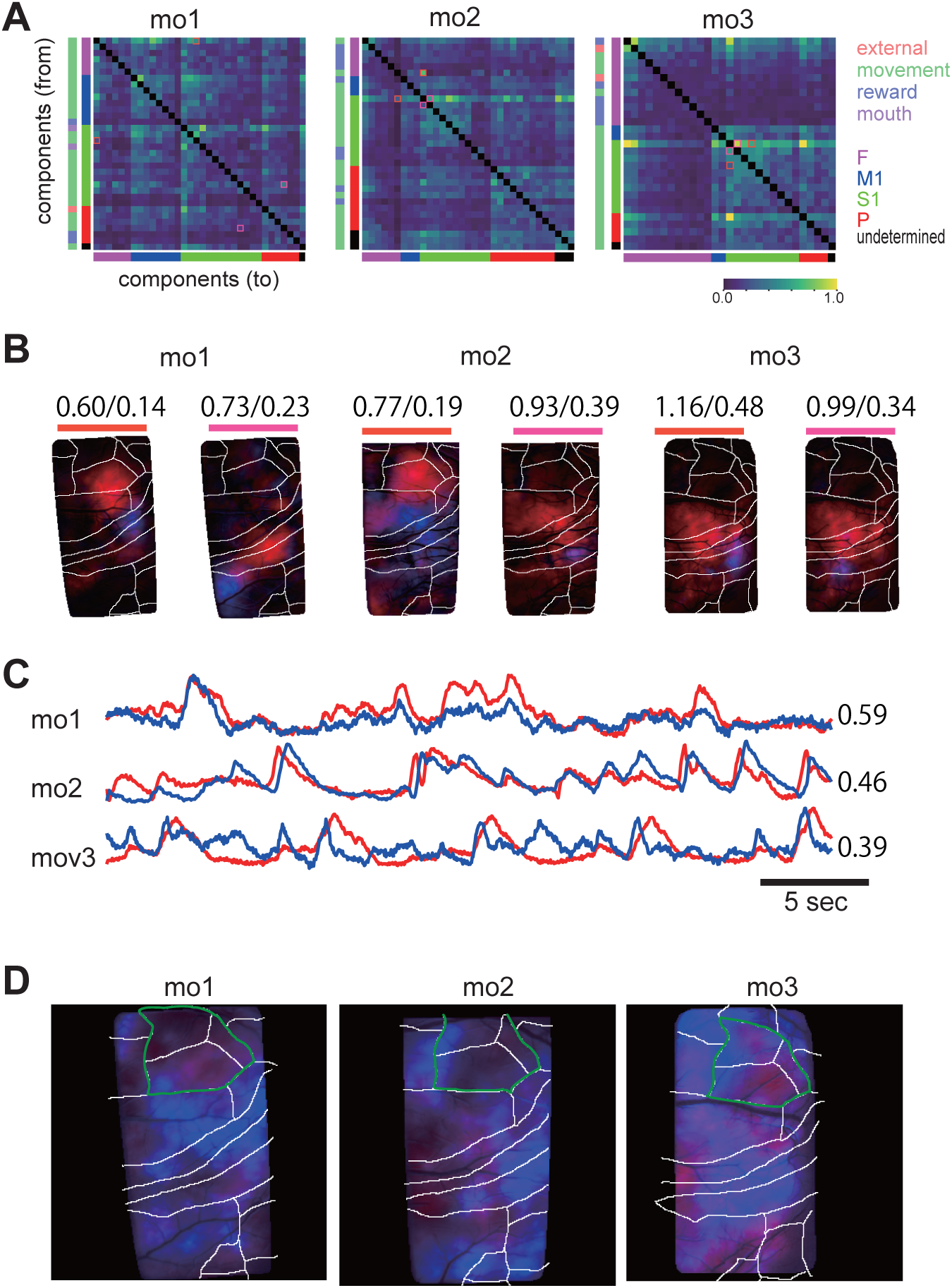
Embedding entropy analysis A: Mean EE for each monkey. The values are ordered by the mean EE values within each area (F, M1, S1, P, and undetermined). The coloured vertical bars indicate the dominant explaining variable category identified in linear regression in Figure 4 (external, movement, reward, and mouth). Red and pink rectangles indicate the two activity component pairs from each monkey that showed a large difference between the EE values in opposite directions. B: Footprints of the activity component pairs indicated by the red and pink rectangles in A. The two numbers at the top are the EE values of the red to blue and blue to red activity components, respectively. C: Example time courses of the activity components shown in B (pink pairs). The right-most numbers are Pearson’s correlation coefficients between the two traces. D: Asymmetry of causal interactions. Activity components were categorized, whether ‘from dominant’ (source) or ‘to dominant’ (target), based on the total EE value from other areas and those to other areas. The from-dominant activity component footprints are coloured red, and the to-dominant activity components are coloured blue. The subareas inside the green lines include 6DC, 6DR, and 8c.

Although matrix is generally symmetric, some pairs showed relatively large differences depending on the prediction direction. Figure 5B shows representative activity component pairs with large EE differences, depending on the inference direction. Despite low correlation coefficients, some activity component pairs exhibited high EE values (Figure 5C; traces for mo3 showed a correlation coefficient of 0.39). The mean EE values were projected onto cortical footprints to examine whether there was a dominance bias in the causal relationships (Figure 5D). The premotor area roughly corresponded to areas 6DC, 6DR, and 8c, which tended to be source-dominant (reddish) in all three monkeys. The motor and somatosensory areas, in contrast, showed both source-dominant and target-dominant areas, and these patterns were not consistent between monkeys. Similarly, the parietal area showed both source- and target-dominant areas.

To evaluate the network property, we created a reciprocal network with activity components as nodes and EE values as edges and examined several network metrics. Supplementary information S5 shows a comparison of these metrics (degree in, degree out, PageRank, and eigencentrality) across the activity components. One or two activity components in premotor area showed much higher values than other activity components in the same area, and their footprints were found in areas 6M or 6Dc. In other areas, activity components in 3a and PE exhibited relatively high values.

### Change of EE values across sessions

Next, we examined changes in EE values across sessions and found the increase trend of EE values across sessions for all monkeys (Figure 6A). To evaluate this observation, we calculated the slopes of the EE values of each activity component pair across sessions using linear fitting. Figure 6B shows the distribution of the slopes of all the pairs. The distribution was shifted in a positive direction (mean: 0.007, 0.004, and 0.01 for mo1, 2, and 3, respectively), indicating that the EE values increased across sessions in all three monkeys. The areas that showed an increase were diverse and not restricted to a particular area (Figure 6C). A similar analysis of the correlation coefficients or transfer entropy between the activity component time courses did not clearly show an increasing trend (data not shown), indicating the EE’s effectiveness of capturing the relationships.

**Figure 6.**
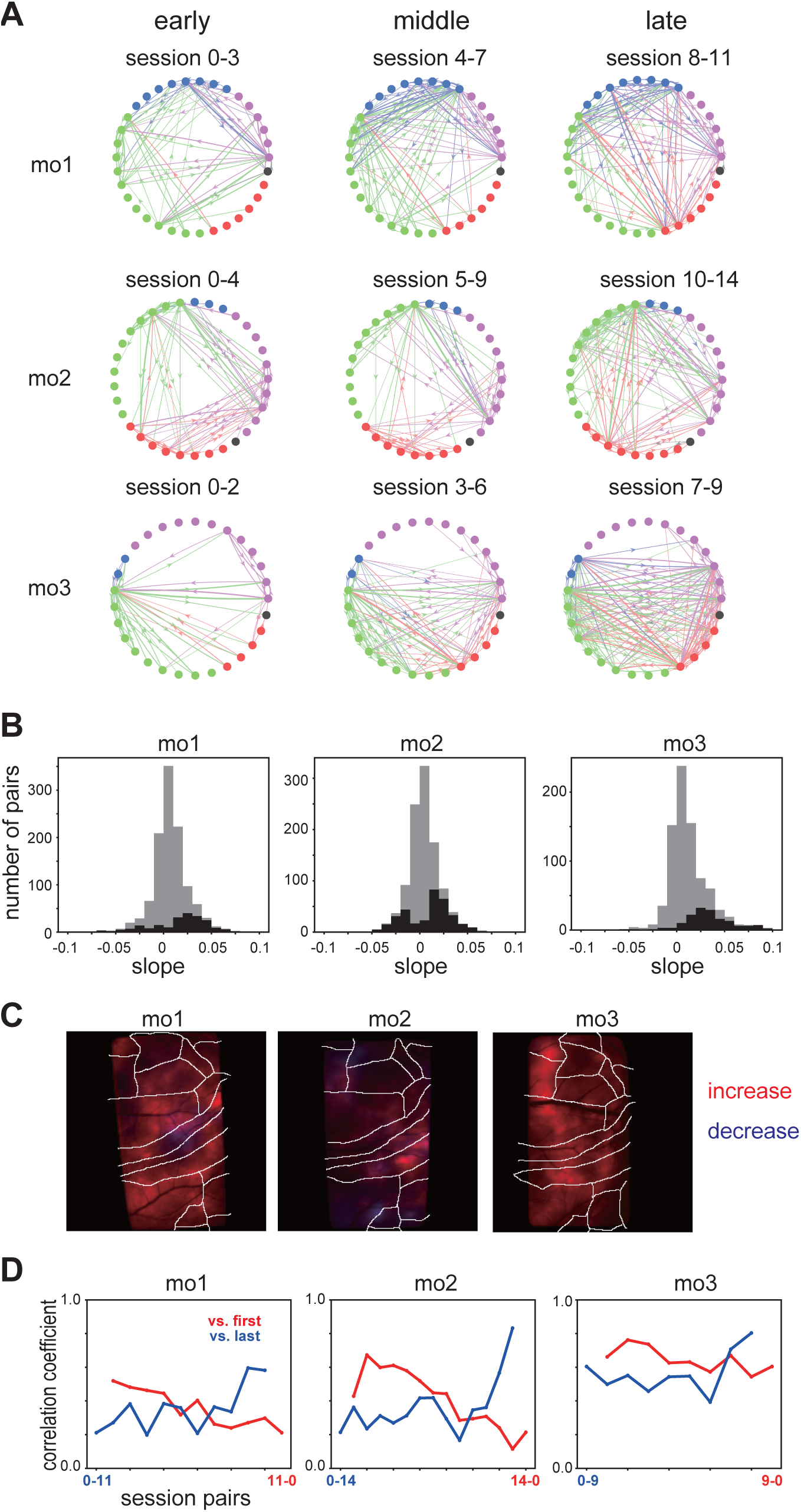
Change of embedding entropy values across sessions A: The network graph shows the EE value change. The EE values were averaged across consective 3-5 sessions for early, middle, and late sessions. Nodes are activity components and the EE values are indicated by the width of edges (thresholded with 0.4) and are colored based on the source nodes. B: Distribution of slope of EE values across sessions; n = 1122, 1056, and 812 for mo1, 2, and 3, respectively (all pairs of activity components). Black histograms correspond to the slopes that showed significant increase or decrease (Wald test, p<0.05). C: Footprints of the activity components that showed a significant slope (p<0.05; Wald test) are projected separate colours for an increase (red) and decrease (blue). D: Correlation coefficients of the EE pattern between the first session and other sessions (red) and between the last session and other sessions (blue).

We examined the consistency of the EE patterns across sessions (Figure 6D). The correlation coefficient between the latter and last sessions was higher than that between the earlier and first sessions, indicating that the pattern stabilized across sessions. To examine whether the observed change in EE values was specific to target-up trials or target-down trials, we stratified the activity component time-courses based on the trial targets and calculated the EE values separately for these two trial types (Figure 7). We then examined the slopes of the EEup and EEdown values across sessions. The slope distribution showed a slight shift in the positive direction in EEup in all three monkeys (mean: 0.002, 0.01, and 0.02 for Monkeys 1, 2, and 3, respectively), and the numbers of significantly positive/negative slopes were 86/83, 176/61, and 126/12 for mo1, 2, and 3, respectively. In contrast, the mean values for the slope of EEdown were −0.002, −0.002, and 0.03 for monkeys 1, 2, and 3, respectively; thus, mo2 and 3 showed a small shift in the decreasing direction. The number of significantly positive/negative slopes was 40/80, 81/88, and 132/2 for mo1, 2, and 3, respectively. These results indicate that the increasing EE values in Figure 6 were mainly the effect of increasing EE values in the up trials in mo1 and 2. Both up- and down-trials contributed to an increase in EE in mo3.

**Figure 7.**
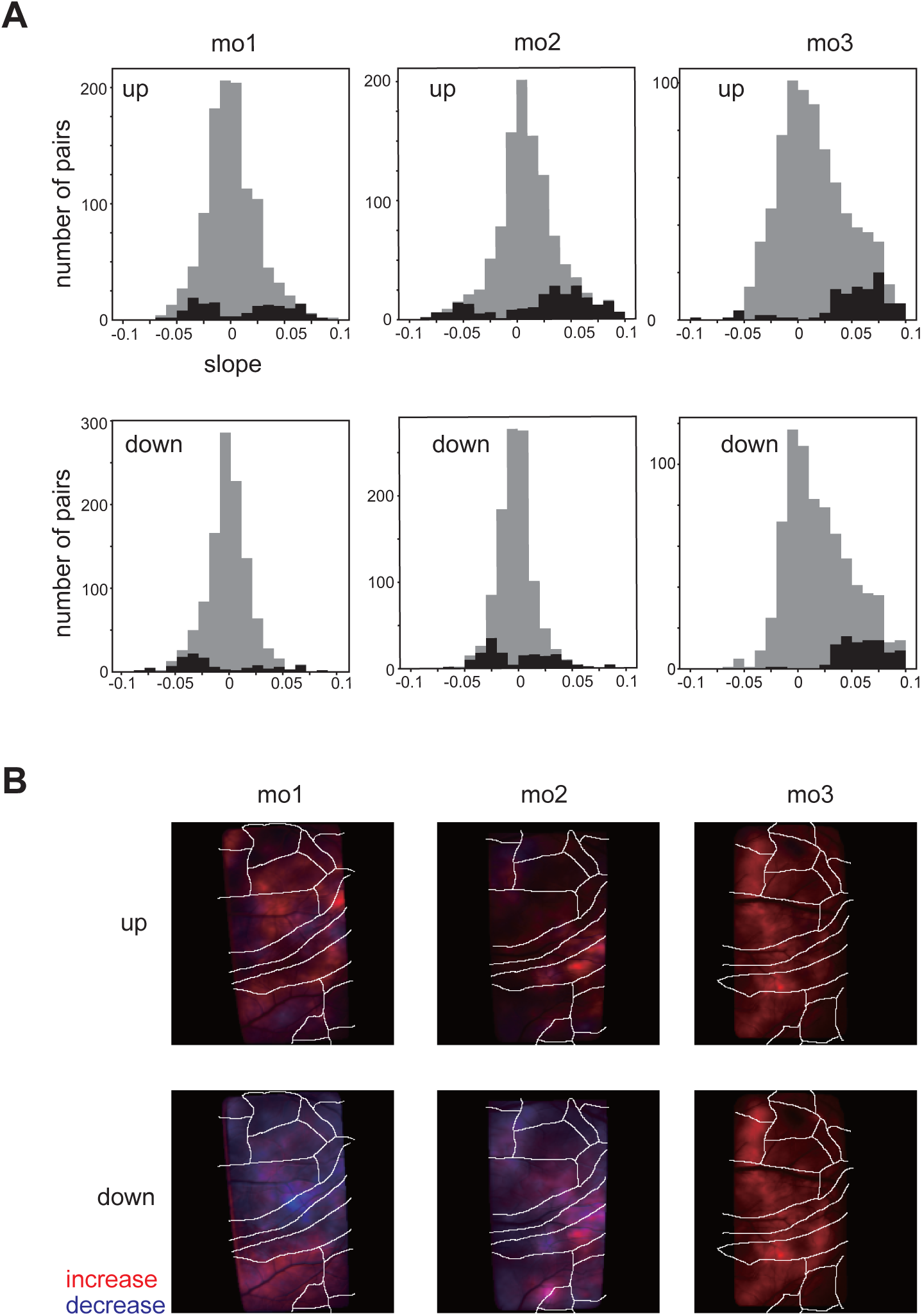
Comparison between EE values calculated based on only up trials (EEup) or only down trials (EEdown). A: Distribution of slopes of EEup (upper panels) and EEdown (lower panels) for all activity component pairs in all sessions. n =1122, 1056, and 812 for mo1, 2, and 3, respectively. Black histograms correspond to slopes that show significant increases or decreases (Wald test, p<0.05). B: Footprints of the activity components that showed a significant slope (p<0.05; Wald test) are projected separately for the increase (red) and decrease (blue) for EEup (upper panels) or EEdown (lower panels).

The results of the EE analysis showed that activity components found in areas 6M, 6DC, 3a, and PE had relatively high mean EE values and high centrality. Activity components found in areas 6DC, 6DR, and 8c were source-dominant. The EE values from these areas were higher than other areas, indicating that these are upstream areas. The EE values increased across sessions, especially in the upward trials. The pattern of the EE network stabilized across sessions. These observations indicate that the neural activity changes widely across brain areas during learning to become more concerted.

## DISCUSSION

In this study, Wide-field calcium imaging and conventional and novel causal analyses focussed on neural dynamics during learning yielded the following results: Cortical activity patterns change from those related-to movement to more consistent with target-related parameters across sessions. The mutual predictability of many activity component pairs, as examined by the EE, increased across sessions, indicating more coordinated activity in later sessions. Moreover, the EE pattern stabilized across sessions. This change was associated with behavioural improvement because it was dominant in the upward trials. These changes cannot be explained by the variance in the decrease in neural signals.

Pan-cortical changes have been observed in the textural discrimination task in mice. Gilad and Helmchen^15^ have reported learning-related cortical changes. Through learning, the mice associated textural stimuli and go/no-go behaviours. They analysed the pre-stimulus and stimulus periods and found that the decrease in activity in several cortical association areas occurred several hundred times before actual learning (they could calculate the learning threshold). Furthermore, they found that enhancement of the task-related cortical activation sequence emerged parallel to increasing task proficiency. In our study, some activity components decreased during learning. We could not determine the learning threshold. Therefore, the timing of the decrease relative to learning was unclear. The stabilization of EE patterns across learning may align with the observation of enhanced task-related cortical activation sequences. Detailed changes in individual areas cannot be compared because of differences in tasks and species. However, the common observation that cortical-wide changes, including increases and decreases in activity across learning, can be captured by wide-field imaging is consistent and noteworthy, indicating a cortical-wide reorganization of activity patterns across learning.

Another study examined the cortical-wide activity change in mice during a leaning lever-pressing task and reported the temporal compression of cortex-wide activity patterns and a decrease in trial variability of activation, which was not explained by variability in behavioural motion^26^. In our study, the change in activity variability did not explain the EE increase (data not shown). It is possible that the tight sequence of activation patterns makes it easier to predict each other, thereby increasing the EE values. Makino et al. observed the emergence of Granger causality from M2 to other cortical areas. We observed an increase in EE for some frontal activity components. Our observation that the EE pattern stabilized across sessions aligns with their results, including widespread increases and decreases in relationships. However, we did not find a single specific area or activity component that showed strong causal relationships with all the other areas, whereas the activity components in A6Dr, A6Dc, and A8c were dominant in the mean EE values. Thus, the observation of the strong influence of M2 in mice^26^ was not entirely consistent with our observations. In our study, multiple patterns of causal relationships were observed, mostly with strong reciprocity. There can be multiple discrepancies, such as differences in task or learning strategies, species, and brain complexity. In addition, this difference may be due to the different methods used to measure causality. Granger causality assumes a linear relationship and separability of dynamics leads causality from its own dynamics, whereas in EE, no such assumptions are made.

In the present study, we observed that some activity components of the somatosensory area, especially Area 3a, showed high centrality, indicating hub-like properties in the network. Motor learning is closely associated with somatosensory feedback^33^. Area 3a processes proprioceptive inputs^31,38,23^, and, together with areas 1 and 2, there are no apparent homologs in the rodent brain^32^. The ICMS for these somatosensory areas in marmosets^6^ and macaques^39, 3^ showed topological maps that are consistent with M1 and Area 3a. In macaque monkeys, corticospinal input has been observed in areas 3a, 3b, and 1/2 ^5^, indicating a strong relationship with motor control. Thus, our observation that Area 3a showed a relatively high centrality is intriguing, as it indicates functional dissociation within the somatosensory area of the cortical networks in primates.

We observed M6 and 6Dr as high-centrality activity components. Some activity components of the parietal area (the PE and PFG) showed high centrality. The parietal and frontal areas are included in the learning network mainly during the early learning phase^8^. Long-train ICMS in the posterior parietal cortex as well as the premotor and motor areas in primates have been demonstrated to elicit complex behaviours, such as reaching, grasping, and defensive movements^20^. The arrangement of modular areas inducing different movements was similar between the motor and parietal areas, suggesting the existence of a parieto-frontal network that controls higher-order movement behaviours^20^. The parietal areas in these studies encompassed a wide range, including the PE, PEc, PGm, 31, V6A, or Areas 5, 7a, 7b, AIP, VIP, and LIP. Among these, only parts of the PE, PF, and PFG were included in the ROI in our study. The major cortical afferents of the PE are the primary somatosensory, motor, and premotor cortices, particularly the regions representing the upper and lower limbs. Furthermore, the PE receives somatosensory inputs from the parietal and cingulate cortices and supplementary motor areas^13^. PE neurons code the direction and depth of forelimb reach^10^. Interestingly, PE (Area 5) neurons have been shown to have motor error information (displacement of the target and hand endpoints) in target-reaching under prism lens conditions^18^. Moreover, neurons in the PFG show mostly hand- and mouth-related activity^14^. Thus, the high centrality observed in the parietal areas in our study might capture the function of the higher-order control of limb behaviour and learning. Recording an extensive parietal network and exploring its functional relationships are intriguing avenues for future research.

To reduce the dimensionality of the imaging data, we used NMF. Other studies have used independent activity component analyses^45^, PCA followed by spatial ICA^26^. The advantages of NMF are its interpretability and ability to demix spatially overlapping signals in a single process. It is also common to delineate areas according to an atlas, particularly in rodent experiments (ex. 48 but see 41). However, this was not performed in the present study. This decision was based on the substantial interindividual variability in primate brains and the fact that we did not image the entire brain, which introduced ambiguity in the map correspondence. In this regard, there is room for improvement in the methodology. In this study, data-driven dimension reduction was employed. In such data-driven methods, the comparison of subjects is generally difficult^40^. Another study on rodents reported variability in activity maps across subjects^15^. Thus, the observation of high inter-subject variability was atypical for our dataset. We observed greater map differences in M1 and S1 than in associated areas. Some variability may arise from differences in detailed functional maps, behavioural strategies, or muscle use. In our study, we were unable to discern the reason.Future work is required to identify the reasons for map variability.

Learning involves processes over multiple timescales^8^. Though the present study focused on changes occurring over several weeks across sessions, it is possible that changes also occurred within individual sessions. Additionally, we assumed the change to be linear in this study; however, nonlinear changes were possible. For example, the correlated activity between M1 neurons increases and decreases within 2–3 weeks of motor learning in rodents^36^. As a unified trend did not emerge consistently across multiple activity components, and because of the complexity of these potential scenarios, addressing these possibilities was beyond the scope of the current study, and these need to be investigated in future work.

## METHODS

### Dataset

The wide-field calcium imaging data used in the analyses of this study are a subset of the data previously reported in Ebina et al^11^. The surgical procedure is described in detail in that study. All experiments were approved by the Animal Experimental Committee of the University of Tokyo and the Animal Care and Use Committees of the RIKEN Center for Brain Science., as previously reported^11^.

### Animals

Three adult common marmosets, *Callithrix jacchus* (two males and one female, weighing 250–350 g), were trained to perform one- and two-target reaching tasks using a two-dimensional manipulandum. All the experiments were approved by the Animal Experimental Committee of the University of Tokyo and the Animal Care and Use Committee of the RIKEN Center for Brain Science.

### Forelimb-reaching task

The details of the marmoset training procedure are provided elsewhere.^12, 11^. The marmosets were seated while wearing a jacket^12^ 17 cm away from a 7-inch LCD monitor. They had already been trained to sit with their heads fixed and perform a target-reaching (pulling) task. They started the two target-reaching tasks after they could perform the one target task (the reward trial rate was ≥70%, with a holding time of ≥400 ms).

The trial began with a holding period wherein the pole was forced to the central position by a spring force of 0.25 N and kept for 1.4–2.1 s. Subsequently, the target (green rectangle) was presented and the spring force was released (reaching period). The animal was rewarded if it moved its pole to match the cursor to the target and maintained it within the assigned area. The target colour changed to white, indicating trial success. Parameters such as duration until time out, holding time, distance between the fixation and target position, and target size changed depending on the performance and training sessions. The final parameter values were 1000–1200 ms for the fixation period, 100 ms for the holding time within the target, 8 × 8 mm for the fixation area size, 16 × 16 mm for the target size, and 14 mm for the distance between the centre of the fixation and the target.

### Wide-field imaging data recordings

As described by Ebina et al^11^, single-photon imaging was conducted using a variable zoom microscope (Axio Zoom.V16; Carl Zeiss, Jena, Germany) equipped with an air objective lens (Plan-NEOFLUAR Z 2.3 ×; numerical aperture 0.5; Carl Zeiss), a fluorescent light source (HXP 200 C; Carl Zeiss), and a filter set (38HE, Carl Zeiss; 470/40 nm excitation filter, 495 nm dichroic mirror, and 525/50 nm emission filter). The imaging FOV was 12.6 x 12.6 mm. The photodetector was a scientific CMOS camera (Sona; Andor Technology) with a resolution of 2048 × 2048 pixels for Marmosets 1 and 2. For Marmoset 3, imaging was conducted using a custom-made zoom-variable microscope equipped with an air objective lens (Plan-NEOFLUAR Z 1.0 ×; numerical aperture 0.25; Carl Zeiss), a fluorescence light source (M470L5; Thorlabs), and a filter set. The imaging FOV size was 14.6 × 14.6 m. The photodetector was a scientific CMOS camera (ORCA-Fusion, Hamamatsu Photonics) with a resolution of 2304 × 2304 pixels. All the images were acquired at a frame rate of 30 Hz.

### Behavioural data recordings

The task events were recorded using LabVIEW (National Instruments) at a sampling rate of 1 kHz. Body and facial movements were recorded using two CMOS cameras (DMK33UP1300; ImagingSource, Taipei, Taiwan). Images were acquired at 320 × 240 pixels at a frame rate of 100 Hz (for marmosets 1 and 2) or 480 × 480 pixels at a frame rate of 50 Hz (for marmoset 3). The recorded movies were used to extract mouth and body movements using Deep Lab Cut (Mathis, 2018; Nath, 2019) available at https://github.com/DeepLabCut/DeepLabCut.

### Pre-processing of wide-field imaging data

Tangential drifts in the imaging were removed using a finite Fourier transform algorithm^9^, and the data were downsampled from 2048 × 2048 or 2304 × 2304 pixels to 256 × 256 pixels. The XY position of frames was registered with the NoRMCorre^37^, available at https://github.com/flatironinstitute/NoRMCorre. Δ*F*/*F* of each pixel of each time *t* was derived by calculating

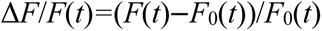

where *F*(*t*) was the fluorescence intensity value at *t*, and *F*_0_(*t*) was the 8th percentile of the *F* value across *t* ±15 s (corresponding to 900 frames).

### Behavioural data analysis

We determined motion onset as the time of onset of a fast large motion immediately before the success of the trial because the monkey sometimes moved its cursor multiple times before success. The mean trajectory deviation across the sessions was fitted using a linear function to calculate the slope. The significance of the slope was tested using Wald’s test.

### NMF

To reduce the dimensionality of the data, we performed MNF for the ΔF/F imaging movies. NMF decomposes the matrix as D ≈ WH, where D is the matrix of n-pixels x m-frames, W is k x n footprints, and H is m x k time series. The number of components or rank (k) was determined by a pilot analysis that examined the data points of one session with different k (5-300). We found that the decrease in reconstruction error became small at k=50 or more (Supplementary information S2-1A). To decompose the concatenated data, we adjusted the movie across sessions using affine transformation because the ΔF/F imaging movies were motion-corrected for each session, and there were slight shifts in the FOV. To find the transformation matrix, we searched for the best-matched blood vessel patterns of two images from different sessions using a scale-invariant feature transform (SIFT). The first 108000 frames of the transformed movies were concatenated across the sessions. The pixels of the concatenated movie were masked to select only the relevant pixels (those inside the FOV that had relevant values across all sessions). We applied NMF to this concatenated movie with L1 regularization on both dimensions of 0.1 and a rank (k) of 50. After decomposition, we manually examined every footprint and removed the components that were highly similar to the blood vessel pattern from further analysis (Supplementary information S2-1B). To calculate the entire range of the time course for all sessions, the matrix of individual sessions were multiplied by the inverse of calculated footprints.

### Encoding analysis

We quantified the time course of each activity component by ordinary linear encoding model.

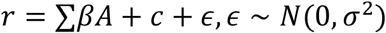

where r, β, c and A denote the time-course of activity component, coefficient, constant, and behavioural parameter values, respectively. For behavioural parameter A, we considered features describing the task variables (target up and down) and behavioural actions (vertical position of the cursor, cursor speed, and mouth-opening angle). The binary variables (target up and down) were multiplied by a Gaussian kernel with a standard deviation of seven frames. To consider the time difference between neural activity and behavioural parameters, we included a lagged version for each parameter (−3.3 to 3.3 sec with an interval of 0.33 s), and the final number of parameters was 189 (nine parameters and 21 lags) for the individual models. The coefficients were determined using LASSO regularization. The hyperparameters for regularization were grid-searched, and the best model evaluated by the coefficient of determination (score) was obtained. The score of each model was calculated based on the fit of 5-fold cross-validation. The models were independently determined for each activity component in each session. To evaluate the importance of each parameter, we created a reduced model in which all regression terms with particular parameters were removed and the same fitting procedure was performed with the full model. The deviance ratio (d-ratio) was calculated as follows:

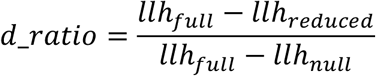

where *llh_full_*, *llh_null_*, and *llh_reduced_* are the log-likelihoods of the full, null, and reduced models, respectively. When the full model performance was very poor and llh-full-llh-null <0, *d_ratio* was set to 0. For the pixel-based regression, the *ΔF/F* of every 8 pixels over FOV were independently used for regression similarly as described above for the regression of NMF components.

We performed the same regression to compare the regression results based on the NMF-extracted activity components and direct regression based on the individual pixel time-course. The detailed results are shown in Figure S4-3.

### Embedding entropy

EE^42^ is a modified causality analysis method based on convergent cross-mapping^43^. Convergence cross-mapping is based on the theory ^44^ of reconstructing attractor manifolds from time-series data. This theory ensures that it is possible to cross-predict or cross-map variables from the same system. We created a dynamic representation of the time series X in an E-dimensional space using a series of E time-lagged values of X to reconstruct an attractor manifold Mx.

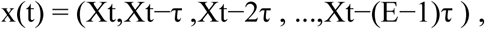

where τ is the time lag. This vector represents the development of X within time t − (E − 1)τ to *t* as a point in E dimensional space. A set of points builds a manifold, Mx. If one of the points in Mx is near another point, the states at these time-points are similar. Thus, the original value of X can be inferred from nearby points in Mx. Cross-mapping between time-series pairs X and Y was performed by predicting Y using Mx or X using My. EEy→x evaluate the inference accuracy by mutual information I between points in Mx and Y instead of correlation coefficient between predicted and actual Y in convergence cross mapping. In our dataset, we calculated the manifold for each activity component time-course and cross-predicted all possible pairs of components. To determine the embedding dimension E and lag τ, we varied E and τ within the range that one point in the manifold is less than 1.5 sec and compared the self-prediction accuracy of the component time series. As the combination of E=9, τ=5 showed the highest accuracy, we used these values for cross prediction. The EE of each component pair in each session was calculated. To calculate EEup and EEdown, the data during the reach time of the up- or down-target trials were separately selected and used to construct the embedded manifold.

## ACKNOWLEDGEMENTS

We are grateful for the help and support provided by the Scientific Computing and Data Analysis section of Research Support Division at OIST and for English Language editing by Editage (www.editage.jp).

## CONFLICT OF INTEREST

The authors declare that this study was conducted in the absence of any commercial or financial relationships that could be construed as potential conflicts of interest.

## FUNDING

The present work was supported by the program for Brain/MINDS from AMED under Grant number P21dm0207001, JP18dm0207027, and JP23wm0625001 to M.M., JP21dm0207085 and JP24wm0625108 to T.E. and M.M., JP21dm0207001h0008 and 20dm0207001h0007 to K.D.

## Notes

### Competing Interest Statement

The authors have declared no competing interest.

