## Supplemental material S1 for "Pan-cortical area sensorimotor network coordination during motor learning of forelimb-reaching task in the marmoset"

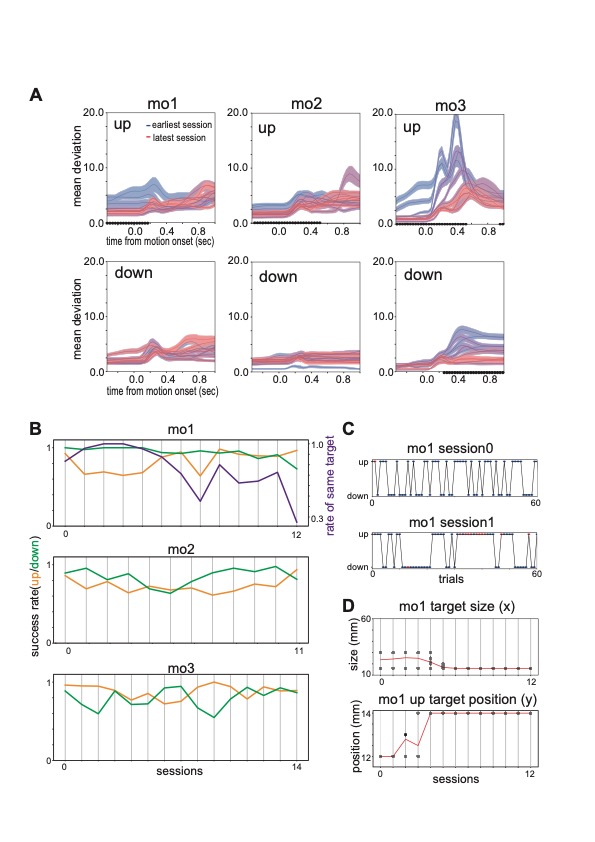

Supplemental figure S1

A: Mean trajectory deviation triggered by motion onset in each session. Colours correspond to different sessions. The coloured areas correspond to the SEM. The black dots on the x-axis are time-points wherein the reduction in mean deviation across sessions was significant (Wald test, p<0.05).

B: Success rate across sessions for individual animals. The rate of the same target just after the failure trials of mo1 is shown as a purple line.

C: Target position of the first 60 trials and the success (blue dots) or failure (red dots) for sessions 0 and 1 of mo1.

D: Target horizontal size and vertical position across session for mo1 (black dots) and their mean across trials (red lines).

We expected an increase in the success rate across sessions in up trials, but the success rate was relatively high in both up and down trials, and an increase was not observed (B). Multiple parameters contribute to the success rate. We explain the reasons using mo1 as an example. In the course of learning, especially early phase of learning, the monkey was exposed to the same target multiple times continuously to enhance learning (C). This setting decreases the success rate. The Success rate (orange line) and rate of the same target (purple line) in (B) are anti-correlated for early sessions. Also, the target's size and position were changed across learning, affecting the success rate (D).
