## Supplemental material S2 for "Pan-cortical area sensorimotor network coordination during motor learning of forelimb-reaching task in the marmoset"

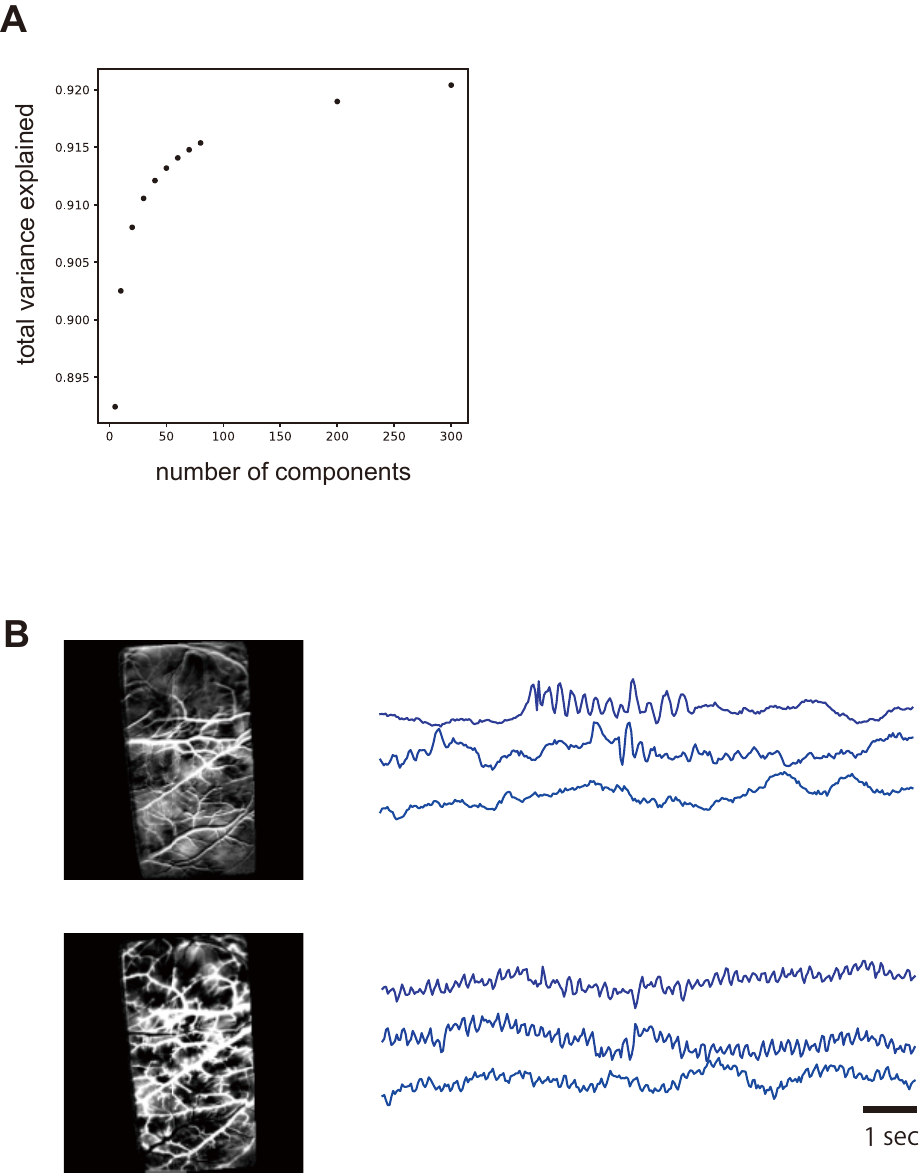

**Supplemental figure S2-1** Details of dimension reduction by NMF.

A: Variance explained across different decomposition ranks for a session of mo1.

B: Examples of component footprints (left) and corresponding time courses from three different sessions (right) of mo1 that showed correlations with blood vessel patterns and were removed from the analysis. Many of the cases, these components contained noise around 6-7Hz, originating from the heartbeat.

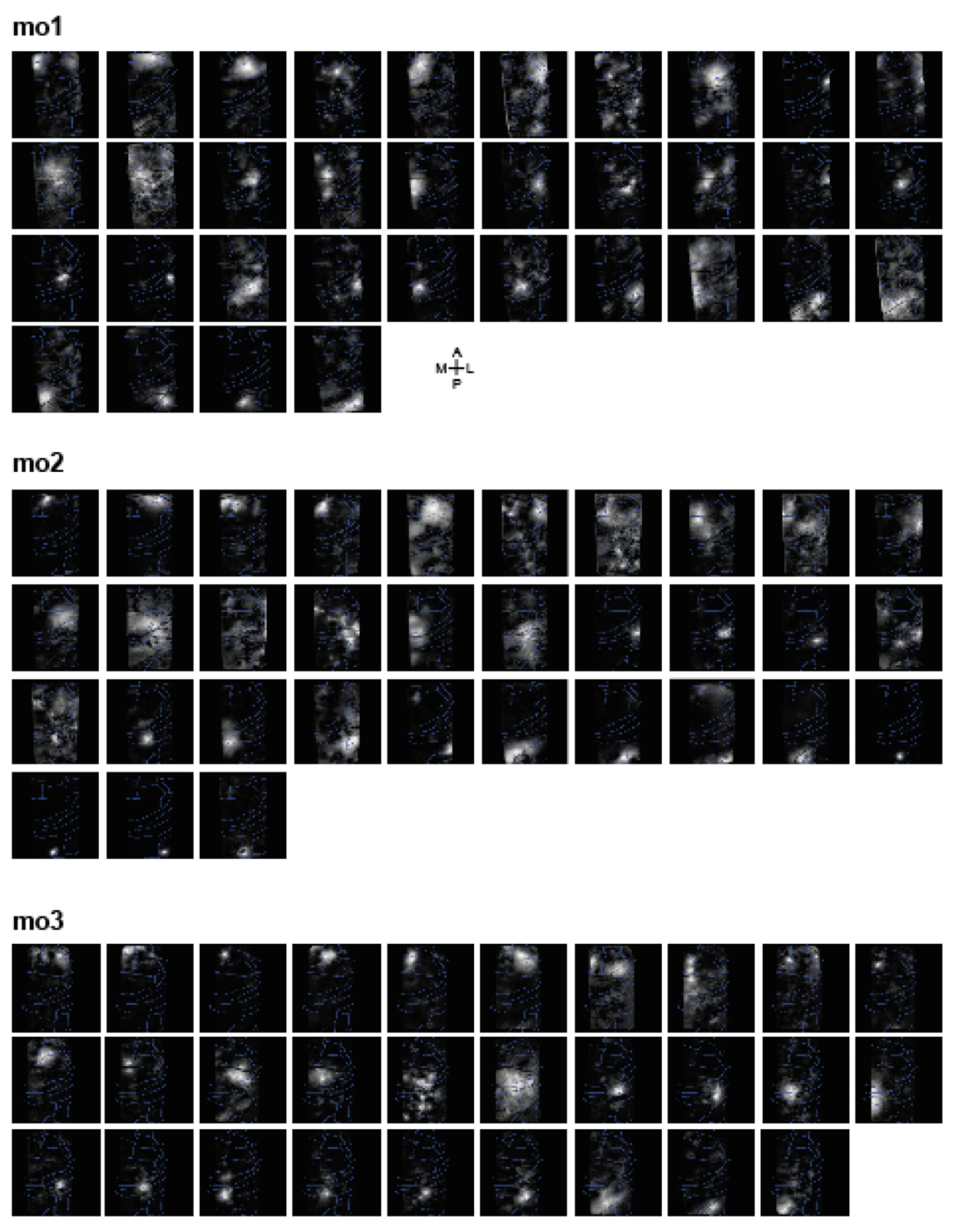

**Supplemental figure S2-2** Footprints of all components included in the analysis for each monkey. The components were aligned from anterior to posterior according to their peak intensity pixel positions. Maximum values were used to scale the intensity of each image for visualization. Blue lines represent putative boundary lines of the brain areas.

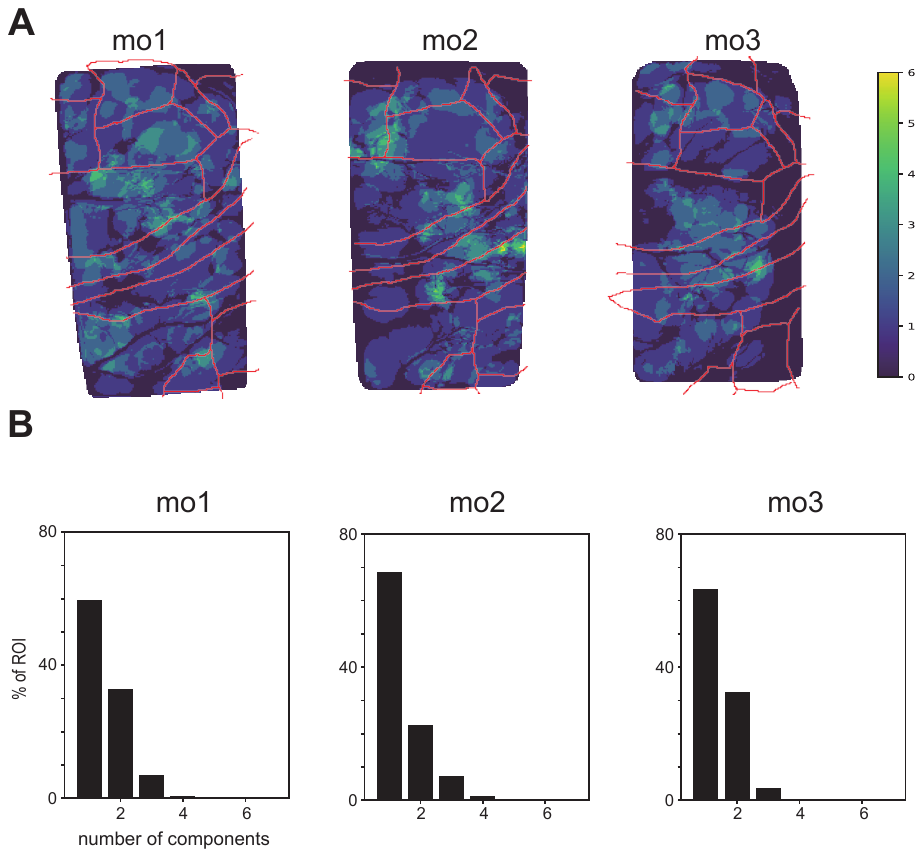

**Supplemental figure S2-3** Overlap of footprints

A: Overlap of footprints based on the 50% peak value of each component. The color corresponds to the number of overlaps.

B: Distribution of the number of components as area % of ROI. One in x axis indicate the area only one component covers the area.
