## Supplemental material S4 for "Pan-cortical area sensorimotor network coordination during motor learning of forelimb-reaching task in the marmoset"

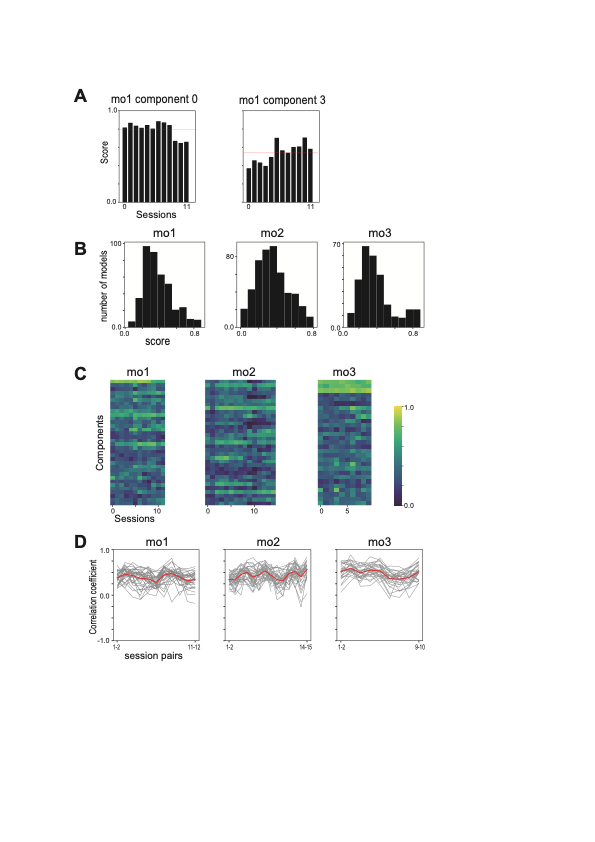

**Supplemental figure S4-1**: A: Explained variance (score) across sessions for the example component in Fig4. The red line represents the mean across the sessions.

B: Distribution of explained variance (score) of all sessions of all components (n=511, 528, and 406 for mo1, 2, and 3, respectively).

C: Explained variance (score) across sessions across components for each monkey.

D: Rank correlation coefficients of the model coefficients (β) between consecutive sessions. The red traces represent the means across the components.

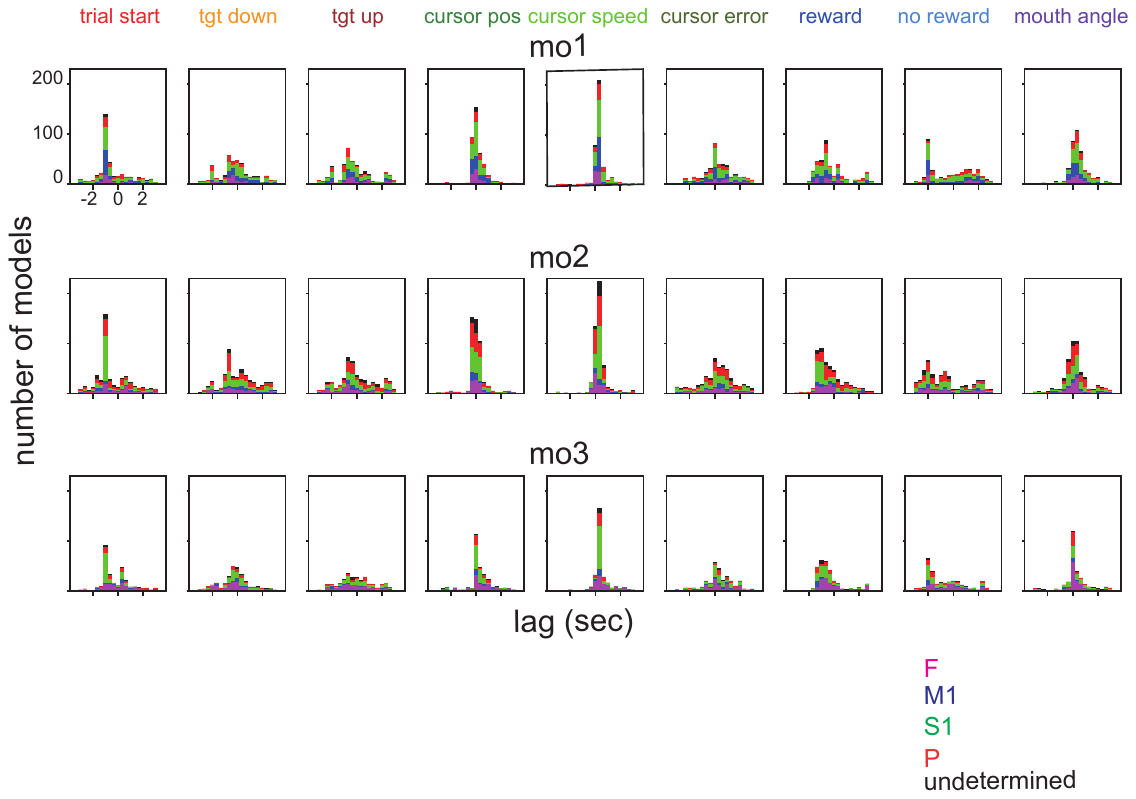

Supplemental figure S4-2: Lag between neural activity and behavior identified by linear regression.

The distribution of peak lag time for individual exogenous variables for each animal is shown. The colours (magenta, blue, green, red, and black) of the bars correspond to the areas (F, M1, S1, P, and undetermined).

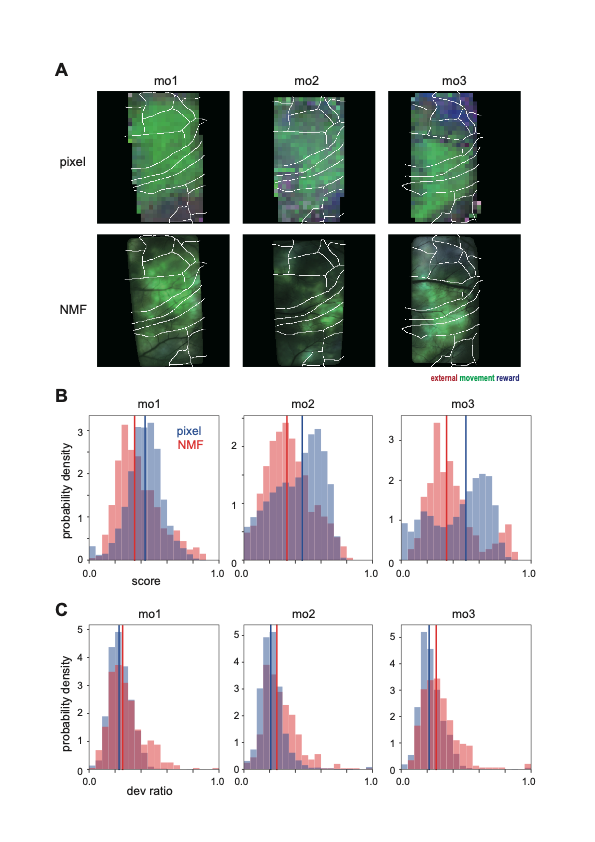

Supplemental figure S4-3 Comparison between pixel-based direct regression and regression of NMF components.

A: The upper row shows the deviance ratio projected onto the cortex. The colors correspond to the behavioral parameter categories (red: movement, green: external, and blue: reward). The lower rows are the same as the upper rows but for the NMF-based regression (They are the same as Figure 4D). B: Comparison of the score distribution between pixel-based (blue) and NMF-based (red) regressions. The distribution was converted to a probability density as the number of examined pixels and NMF components differed. Coloured vertical lines indicate the mean of distribution. C: Comparison of the deviance-ratio distribution between pixel- (blue) and NMF-based (red) regressions. The conventions are the same as those in B.

**Comparison with model fit based on pixel-based time series**

We performed the same regression to compare the regression results based on the NMF-extracted components and direct regression based on the individual pixel time-course. Supplemental Figure S4-3A shows deviance ratios projected onto the cortex, with similar general patterns for the pixel- and NMF-based regressions. The mean scores were higher in the pixel-based regression, though the distribution’s tail extended towards 1 in the NMF-based regression; thus, a small number of components showed high scores in the NFM-based regression (Figure S4-3B). Moreover, the maximum deviance ratio (across parameters) for the NMF-based regression distribution was skewed towards 1 (Figure S4-3C). These observations indicate that a few NMF components were highly dependent on one of the parameter categories and were demultiplexed when compared with the results of pixel-based analysis.
