## Supplemental material S5 for "Pan-cortical area sensorimotor network coordination during motor learning of forelimb-reaching task in the marmoset"

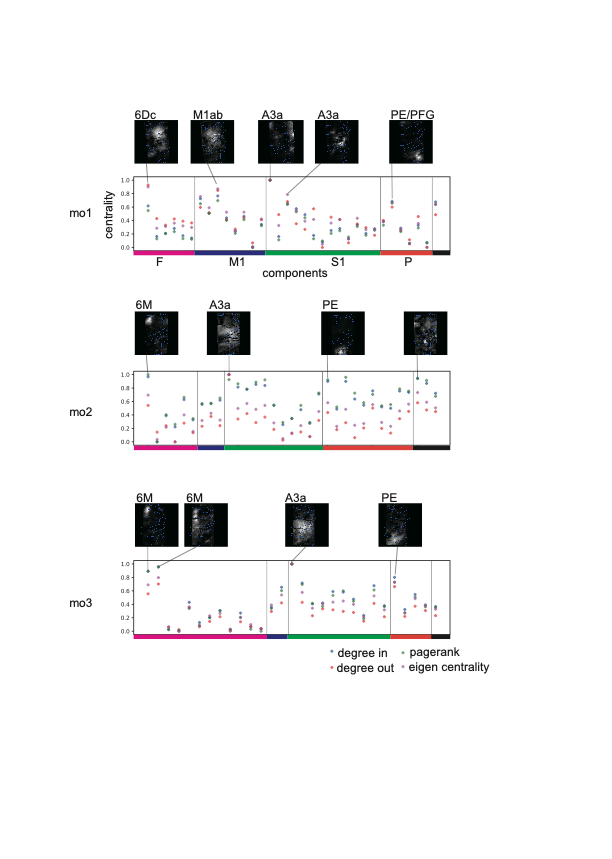

Supplemental figure S5. Comparison of several network metrics between components.

The degree in (blue), degree out (red), PageRank (green), and eigencentrality (magenta) are shown for each component ordered by area. Values are the means across sessions and scaled individually for each metric across the components for each monkey. The insets show the footprints of the example components with high values.
